# Functional and structural analysis of a cyclization domain in a cyclic β-1,2-glucan synthase

**DOI:** 10.1101/2023.10.30.564714

**Authors:** Nobukiyo Tanaka, Ryotaro Saito, Kaito Kobayashi, Hiroyuki Nakai, Shogo Kamo, Kouji Kuramochi, Hayao Taguchi, Masahiro Nakajima, Tomoko Masaike

## Abstract

Cyclic β-1,2-glucan synthase (CGS) is a key enzyme in production of cyclic β-1,2-glucans (CβGs) which are involved in bacterial infection or symbiosis to host organisms. Nevertheless, a mechanism of cyclization, the final step in the CGS reaction, has not been fully understood. Here we performed functional and structural analyses of the cyclization domain of CGS alone from *Thermoanaerobacter italicus* (TiCGS_Cy_). We first found that β-glucosidase-resistant compounds are produced by TiCGS_Cy_ with linear β-1,2-glucans as substrates. The ^1^H-NMR analysis revealed that these products are CβGs. Next, action pattern analyses using β-1,2-glucooligosaccharides revealed a unique reaction pattern: exclusive transglycosylation without hydrolysis and a hexasaccharide as the minimum length of the substrate. Nevertheless, longer substrate β-1,2-glucooligosaccharides are preferred, being consistent with the fact that CGSs generally produce CβGs with degrees of polymerization of around 20. Finally, the overall structure of the cyclization domain of TiCGS_Cy_ was found to be similar to those of β-1,2-glucanases in phylogenetically different groups. Meanwhile, the identified catalytic residues indicated clear difference in the reaction pathways between these enzymes. Therefore, we propose a novel reaction mechanism of TiCGS_Cy_. Thus, we propose that the present group of CGSs should be defined as a new glycoside hydrolase family, GHxxx.

**Statements and Declarations:** Competing Interests: The authors declare no competing interests.

**Key Points:** It was clearly evidenced that cyclization domain alone produces cyclic β-1,2-glucans.

The domain exclusively catalyzes transglycosylation without hydrolysis.

The present catalytic domain should be defined as a new glycoside hydrolase family.

## Introduction

β-1,2-glucan is a polysaccharide comprising glucose units linked by β-1,2-glucosidic bonds. In nature, cyclic forms are mainly found in bacteria such as *Agrobacterium tumefacies*, *Brucella abortus* and *Ensifer meliloti* (formerly *Rhizobium meliloti* and *Sinorhizobium meliloti*) (Dell et al. 1983; Koizumi et al. 1984; Bundle et al. 1988). Cyclic β-1,2-glucans (CβGs) play important roles in interactions between organisms such as infection of *B. abortus* and symbiosis of *E. meliloti* (Breedveld and Miller 1994; Haag et al. 2010; Bontemps-Gallo et al. 2017). Genes encoding enzymes responsible for CβG biosynthesis were identified from three microorganisms independently. These genes were found to encode cyclic β-1,2-glucan synthases (CGSs) homologous to each other (Zorreguieta et al. 1985; Zorreguieta and Ugalde 1986; Castro et al. 1996; Iannino et al. 1998). Among them, CGS from *B. abortus* (BaCGS) produces CβGs with degrees of polymerization (DPs) around 20 and is most extensively characterized. BaCGS is composed of three regions responsible for the following steps: initiation (covalent bonding of glucose to CGS), elongation of linear β-1,2-glucan (LβGs) chains, adjustment of chain lengths of the glucans, and cyclization of the glucans by transglycosylation (Ciocchini et al. 2007). The initiation and the elongation steps are carried out in the N-terminal domain classified into glycosyltransferase (GT) family 84 based on amino acid sequence homology by Carbohydrate-Active enZYmes Database (CAZy) (Coutinho et al. 2003; Drula et al. 2022). The C-terminal glycoside hydrolase (GH) family 94 domain adjusts the chain lengths of the elongated glucans. Although the domain in the middle region is known to be involved in cyclization of the linear glucans of the optimum length (Hereafter, this domain is called cyclization domain), a detailed reaction mechanism has not been unveiled.

Apart from the context of synthetic enzymes described above, β-1,2-glucan-degrading enzymes have been investigated (Nakajima 2023). Owing to establishment of a large-scale preparation of LβGs by using a 1,2-β-oligoglucan phosphorylase (SOGP) found in 2014 (Nakajima et al. 2014; Abe et al. 2015), prokaryotic and eukaryotic *endo*-β-1,2-glucanases (SGLs) that produce β-1,2-glucooligosaccharides (Sop_n_s, n is DP) from β-1,2-glucans were explored. Consequently, a bacterial SGL from *Chitinophaga pinensis* (CpSGL) and a fungal SGL from *Talaromyces funiculosus* (TfSGL) were sequentially identified for the first time, leading to creation of new families (GH144 and GH162), respectively (Abe et al. 2017; Tanaka et al. 2019). Subsequently, functions and structures of several β-1,2-glucan-associated enzymes including the ones that are given new EC numbers were reported (Nakajima et al. 2016; Nakajima et al. 2017; Shimizu et al. 2018; Kobayashi et al. 2022). In addition, another SGL which neither belongs to GH144 nor GH162 has been identified from *Escherichia coli* recently, leading to foundation of a new family, GH186 (Motouchi et al. 2023).

Interestingly, both CpSGL and TfSGL possess single (α/α)_6_-domains with similar overall structures although they belong to different families. Therefore, PSI-BLAST search was performed using CpSGL and TfSGL as queries to find further homologs with evolutional relationships. As a result, the cyclization domains of CGSs (CGS_Cy_s) came up although the domains do not belong to any GH families. The acid catalyst of TfSGL, one of the two catalytic residues, are highly conserved among the cyclization domains, which may suggest that GH162 and the group of CGSs partly share a common reaction mechanism. However, TfSGL and CGS_Cy_ are intrinsically different in terms of reaction mechanisms. TfSGL follows the anomer-inverting mechanism in which anomer of substrates changes when products are released, while CGS follows the anomer-retaining mechanism in which substrates and products share the same anomer (see https://www.cazypedia.org/index.php/Glycoside_hydrolases#Mechanistic_classification in detail) (Davies and Henrissat 1995). With these facts, we presume that CGS_Cy_ follows a novel reaction mechanism. In this study, we subcloned a region encoding cyclization domain alone from CGS of *Thermoanaerobacter italicus*, a thermophilic bacterium, (TiCGS) and explored biochemical functions and tertiary structure of the cyclization domain.

## Materials and Methods

### Materials

The genomic DNA of *T. italicus* (DSM9252) was purchased from the National Institute of Technology and Evaluation (NITE, Tokyo, Japan). LβGs with the average DP of 77 (unless otherwise described, average DP of the β-1,2-glucans is 77.) and Sop_n_s with DP of 2–10 were prepared using SOGP from *Listeria innocua* and CpSGL as described previously (Nakajima et al. 2014; Abe et al. 2015; Abe et al. 2017). CβG with DP of 17–24 was kindly donated by Dr. M. Hisamatsu of Mie University (Hisamatsu *et al*. 1984). Laminarin and carboxymethyl (CM)- cellulose were purchased from Sigma-Aldrich (MO, USA). CM-pachyman, CM-curdlan, lichenan, amarind xyloglucan, arabinogalactan, arabinan and polygalacturonic acid were purchased from Megazyme (Wicklow, Ireland)..

### Cloning, Expression, and Purification of TiCGS_Cy_

A middle region (1005–1591 a.a., TiCGS_Cy_) of TiCGS (KEGG locus, Thit_1831) was used for cloning (see Results for details). A gene region encoding TiCGS_Cy_ was amplified by PCR using PrimeSTAR Max DNA Polymerase (Takara Bio, Shiga, Japan) according to the manufacturer’s instructions.

The PCR was performed using cloning primers and the genomic DNA pool of *T. italicus* as a template. The amplified product digested with NdeI and XhoI (Takara Bio) was ligated into the pET30a vector (Merck, NJ, USA) which was digested with the same restriction enzymes. *E. coli* Rosetta2 (DE3) (Merck) was transformed using the constructed plasmid and cultured at 37°C in LB medium containing 30 μg/ml kanamycin and 34 μg/ml chloramphenicol. After the optical density of the culture at 660 nm reached 0.8, protein expression was induced using 0.1 mM isopropyl-β-D-1-thiogalactopyranoside at 20°C overnight. The harvested cells were lysed by sonication in 50 mM Tris-HCl buffer (pH 8.0) containing 150 mM NaCl. The supernatant was collected after centrifugation at 27,700 × *g*. Then the supernatant was filtrated with a 0.45 µm filter (Sartorius, Germany). The sample was loaded onto a HisTrap FF crude column (5 mL; Cytiva, MA, USA) equilibrated with 50 mM Tris-HCl buffer (pH 8.0) containing 150 mM NaCl (buffer A). After unbound proteins were washed out using the same buffer containing 20 mM imidazole, TiCGS_Cy_ was eluted using a linear imidazole concentration gradient (20–300 mM) in buffer A. 2M ammonium sulfate solution containing 100 mM sodium acetate buffer (pH 5.0) was added to the collected sample to obtain 1 M ammonium sulfate concentration. After unbound proteins was washed out using the 1 M ammonium sulfate containing 100 mM sodium acetate buffer (pH 5.0), TiCGS_Cy_ was eluted using a linear ammonium sulfate concentration gradient (1– 0 M) in 100 mM sodium acetate buffer (pH 5.0). The enzyme solution was exchanged with 5 mM sodium acetate buffer (pH 5.0) using Amicon Ultra 10,000 molecular weight cut-off (Merck). The absorbance of the sample at 280 nm was measured using a spectrophotometer V-650 (Jasco, Tokyo, Japan), and the concentration of the enzyme was determined spectrophotometrically at 280 nm using the theoretical molecular mass of TiCGS_Cy_ (69,508 Da) and a molar extinction coefficient of 87,210 M^−1^·cm^−1^ calculated based on Pace *et al*.(Pace *et al*. 1995).

### Size-exclusion chromatography

The enzyme solution concentrated with Amicon Ultra 10,000 molecular weight cut-off to 0.5 mg/ml (500 μl) was loaded onto a Superdex^TM^ 200GL column (24 ml; Cytiva) equilibrated with 50 mM Tris-HCl buffer (pH 8.0) containing 150 mM NaCl, and then the target enzyme was eluted with the same buffer. Purification step by size-exclusion chromatography was carried out using an AKTA prime plus chromatography system (Cytiva). Ovalbumin (44 kDa), conalbumin (75 kDa), aldolase (158 kDa), ferritin (440 kDa), and thyroglobulin (669 kDa) (Cytiva) were used as molecular weight markers. Blue dextran 2000 (2,000 kDa) was used to determine the void volume of the column. The molecular weight of TiCGS_Cy_ was calculated using Equation 1,

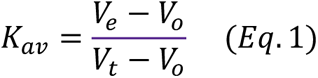

where *K*_av_ is the gel-phase distribution coefficient; *V*_e_ is the volume required to elute each protein; *V*_o_ is the volume required to elute blue dextran 2000; and *V*_t_ is the bed volume of the column.

### Analysis of the cyclization activity of TiCGS_Cy_

The enzymatic reaction of TiCGS_Cy_ on LβG was performed in 20 mM sodium acetate buffer (pH 5.0) containing 1 mg/mL of TiCGS_Cy_ and 0.4% LβG with an average DP of 77 at 30°C for one hour. After a heat treatment at 100°C for 5 minutes, the sample (20 µL) was mixed with 20 µL of 0.1 mg/mL β-glucosidase (BGL) from *Bacteroides thetaiotaomicron* (Ishiguro *et al*. 2017) in 100 mM sodium acetate buffer (pH 5.0) and incubated at 40°C for 30 min or 60 min. After a heat treatment at 100°C for 5 minutes, the sample (20 µL) was mixed with 20 µL of 0.2 mg/mL TfSGL in 100 mM sodium acetate buffer (pH 5.0). The reaction mixture was incubated at 30°C for an hour. Each reaction mixture was analyzed by thin layer chromatography (TLC).

### TLC analysis

The reaction mixtures (0.5, 1 or 2 µL) were spotted onto TLC Silica Gel 60 F_254_ plates (Merck). As for analysis of cyclization activity, the plates were developed with 70% acetonitrile. In the case of glucose and Sop_2-5_ produced by TiCGS_Cy_, the plates were developed with 75% acetonitrile. In the case Sop_6-10_, the plates were developed twice with the solution (1-butanol: acetic acid: deionized water =2:1:1). The plates were then soaked in a 5% sulfuric acid/ethanol solution (w/v) and heated in an oven until the spots were clearly visualized.

The enzymatic reactions of TiCGS_Cy_ on each substrate (0.03% arabinogalactan, 0.2% CM-pachyman, 0.2% laminarin, 0.2% CM-cellulose, 0.2% CM-curdlan, 0.2% arabinan, 0.2% polygalacturonic acid, 0.2% LβG or 0.2% Tamarind xyloglucan, 5 mM glucose and Sop_2–10_) were carried out in 100 mM sodium acetate buffer (pH 5.0) containing 1 mg/mL of TiCGS_Cy_ at 30°C. After heat treatment at 100°C for 5 minutes, the reaction products were detected by TLC.

### NMR analysis

To collect the cyclic products of TiCGS_Cy_ when using LβG as a substrate, the enzymatic reaction was performed at 30°C for 42 hours in 100 mM sodium acetate buffer (pH 5.0) containing 2 mg/mL of TiCGS_Cy_ and 5% LβG with an average DP of 100. After a heat treatment at 100°C for 10 minutes, the supernatant (4 mL) was collected after centrifugation at 4,427 × *g*. Then the solution was mixed with 250 µL of 2.5 mg/mL BGL in 100 mM sodium acetate buffer (pH 5.0) and incubated at 30°C for 27 hours so that the spot of the polysaccharides no longer changes. After a heat treatment at 100°C for 10 minutes, the sample was centrifugated at 4,427 × *g*. Then the supernatant was filtrated with a 0.45 µm filter (Sartorius). The sample was fractionated by size-exclusion chromatography and the fractions containing the target compound was freeze-dried using the FDU-2100 (EYELA, Tokyo, Japan). The resultant powder was dissolved in D_2_O, and acetone was added as a standard for calibration of chemical shifts. The chemical shifts were recorded relative to the signal of the methyl group of the internal standard acetone (2.22 ppm). As a reference, CβG donated by Dr. Hisamatsu (Hisamatsu et. al. 1984) was also dissolved in the same solvent. ^1^H-NMR spectra were recorded using a Bruker Advance 400 spectrometer (Bruker BioSpin).

### X-ray Crystallography

The enzyme solution was concentrated to 17.6 mg/ml. The initial screening of TiCGS_Cy_ crystallization was performed using MembFac HT (Hampton research, CA, USA). The crystal for data collection was obtained by incubation of the mixture of 17.6 mg/ml TiCGS_Cy_ (2 μl) and a reservoir solution (2 μl, containing 0.1 M sodium cacodylate, 1.3 M sodium acetate (pH 6.5)) at 20 °C for one month. The crystal was soaked in the reservoir solution supplemented with 25% (w/v) glycerol for cryoprotection and kept at 100 K in a nitrogen-gas stream during data collection. The X-ray diffraction data was collected on a beamline (BL-5A) at Photon Factory (Tsukuba, Japan). The TiCGS_Cy_ structure was determined by molecular replacement using the predicted TiCGS_Cy_ structure by AlphaFold2 (Jumper *et al*. 2021) as a model structure. The molecular replacement, auto model building, and refinement were performed using the MOLREP program (Vagin and Teplyakov 2010), REFMAC5 program (Murshudov et al. 1997) and Coot program (Emsley and Cowtan 2004), respectively. A structural homology search was performed with the DALI server (Holm 2020). The secondary structure was assigned with the DSSP program (Touw et al. 2015), and the multiple amino acid alignment and the structure-based alignment, including that of the secondary structures, were visualized using the ESPript 3.0 server (http://espript.ibcp.fr/ESPript/ESPript/) (Robert and Gouet 2014). The overall structures of TiCGS_Cy_, TfSGL and CpSGL were superimposed using the PDBeFold server (https://www.ebi.ac.uk/msd-srv/ssm/ssmcite.html) (Krissinel and Henrick 2004). All the structures in the figures were designed with the PyMOL program.

### Mutational analysis

The plasmids of TiCGS_Cy_ mutants (E1442Q, E1442A and E1356A) were constructed using a PrimeSTAR mutagenesis basal kit (Takara Bio) according to the manufacturer’s instructions. PCRs were performed using appropriate primer pairs (Table S2) and the template TiCGS_Cy_ plasmid. The transformation to *E. coli*, the expression, and purification of TiCGS_Cy_ mutants were performed in the same manner as that for the wild-type TiCGS_Cy_. The enzymatic reactions were performed basically in the same manner as in the detection of cyclization activity of TiCGS_Cy_.

## Results

### Purification of cyclization domain of CGS from *T. italicus* (TiCGS_Cy_)

In TiCGS, domain configuration and biochemical function including cyclization activity are totally unknown. Therefore, multiple amino acid alignment was performed using CGS homologs including TiCGS and BaCGS (Fig. S1). The region of cyclization domain was expected to be residues 1005–1591 a.a. based on the minimum region that retains cyclization activity in BaCGS (Ciocchini et al. 2006; Guidolin et al. 2009). In addition, all transmembrane regions are within residues 1–1004 a.a. according to TMHMM-2.0 server (Krogh et al. 2001). Thus, the region (1005–1591 a.a.) was defined TiCGS_Cy_, and TiCGS_Cy_ fused with histidine-tag at the C-terminus was produced as a recombinant protein. The recombinant TiCGS_Cy_ (hereafter simply called TiCGS_Cy_) was purified by nickel affinity chromatography and hydrophobic chromatography, with which we obtained highly purified TiCGS_Cy_ that migrated as a single band at approximately 70 kDa in the SDS-PAGE analysis. It is consistent with a theoretical molecular mass of TiCGS_Cy_ (69.5 kDa).

### Size-exclusion chromatography analysis of TiCGS_Cy_

To investigate quaternary structure of TiCGS_Cy_, size-exclusion chromatography was performed. TiCGS_Cy_ eluted at the retention time corresponding to 60.6 kDa, which is similar to the theoretical molecular mass shown above (Fig. S2). This result indicated that TiCGS_Cy_ exists as a monomer and TiCGS does not form multimer through interactions between the TiCGS_Cy_ domains.

### Cyclization activity of TiCGS_Cy_

To test the ability of purified TiCGS_Cy_ in cyclization of LβG, the reaction products were analyzed by TLC. If TiCGS_Cy_ possesses cyclization activity, both linear and cyclic glucan chains are expected to be produced. After incubation of LβG with TiCGS_Cy_, a broad smear band in DPs smaller than those of the substrate was detected (Fig. 1). Then, the reaction products were subjected to a BGL that act exolytically on the non-reducing end of LβGs (Fig. 1b). Consequently, glucose was produced, but a bit smear band with relatively higher DPs remained on the TLC plate (Fig. 1a). The BGL-treated products were further treated with CpSGL, an endo-type enzyme that produces Sop_2–4_ (Fig. 1b), resulting in disappearance of the smear band and appearance of Sop_2–4_ instead (Fig. 1a). These results indicate that the products that formed the smear band after the BGL treatment were in cyclic forms, and thus TiCGS_Cy_ possesses the cyclization activity. In addition, the smear band after the BGL treatment was detected at the position lower than the spot of the marker CβGs with DP17–24 (M2) in the TLC plate (Fig. 1a), suggesting that DPs of the cyclic products released by TiCGS_Cy_ are higher than 17–24. Furthermore, various polysaccharides were examined as candidate substrates by TLC analysis, but no reaction was detected (Fig. S3). This result suggested that the cyclization activity of TiCGS_Cy_ is highly specific to LβG.

**Figure 1.**
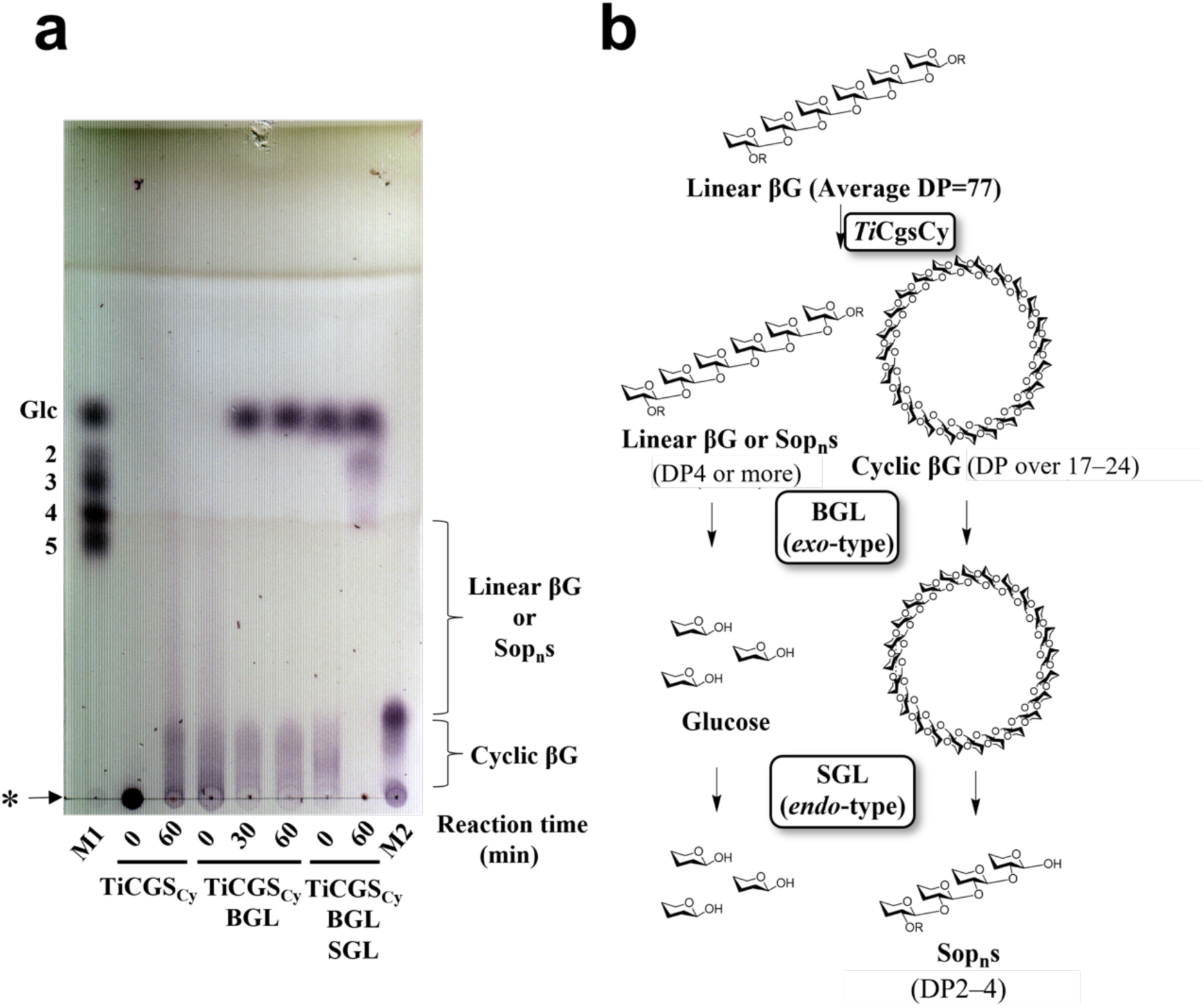
Products from catalysis of LβGs by TiCGS_Cy_. **a**. Detection of the reaction products by TLC analysis. Lane M1, 5 mM glucose and Sop_2–5_. Lane M2, 0.2% CβGs with DP17–24. Each sample (0.5–2 μl) was spotted on the plate. BGL and SGL represent treatment of products with BGL and/or SGL. The asterisk represents the origin of the TLC plate. **b**. The present method to distinguish between cyclic and linear forms of β-1,2-glucans.

### NMR analysis of compounds produced from LβG by TiCGS_Cy_

In order to identify cyclic glucans produced by TiCGS_Cy_, the reaction product from LβG was treated with BGL, and the BGL-resistant products were then purified by size-exclusion chromatography. The chemical shifts of the resultants measured by ^1^H-NMR were almost the same as that of the reference (Fig. S4b) (Hisamatsu et al. 1984; Roset et al. 2006). In addition, chemical shifts derived from H-2 and H-4 at the non-reducing end glucose moiety and H-1 (α-anomer) at the reducing end glucose moiety as in the case of LβG (Nakajima et al. 2014) (Fig. S4c) were not detected (Fig. S4a). These facts indicate that TiCGS_Cy_ produces CβG, and this enzyme follows an anomer-retaining mechanism.

### Action patterns of TiCGS_Cy_

To clarify chain length specificity of substrates, various Sop_n_s were adopted in the experiments. In the case of glucose and Sop_2–5_, no reaction product was detected by TLC analysis (Fig. 2a). On the contrary, Sop_n_s with DPs 4 or higher were produced when Sop_n_s with DPs 6 or higher were applied as substrates. These results indicate that specific substrates of TiCGS_Cy_ in transglycosylation is Sop_n_s with DPs 6 or higher.

**Figure 2.**
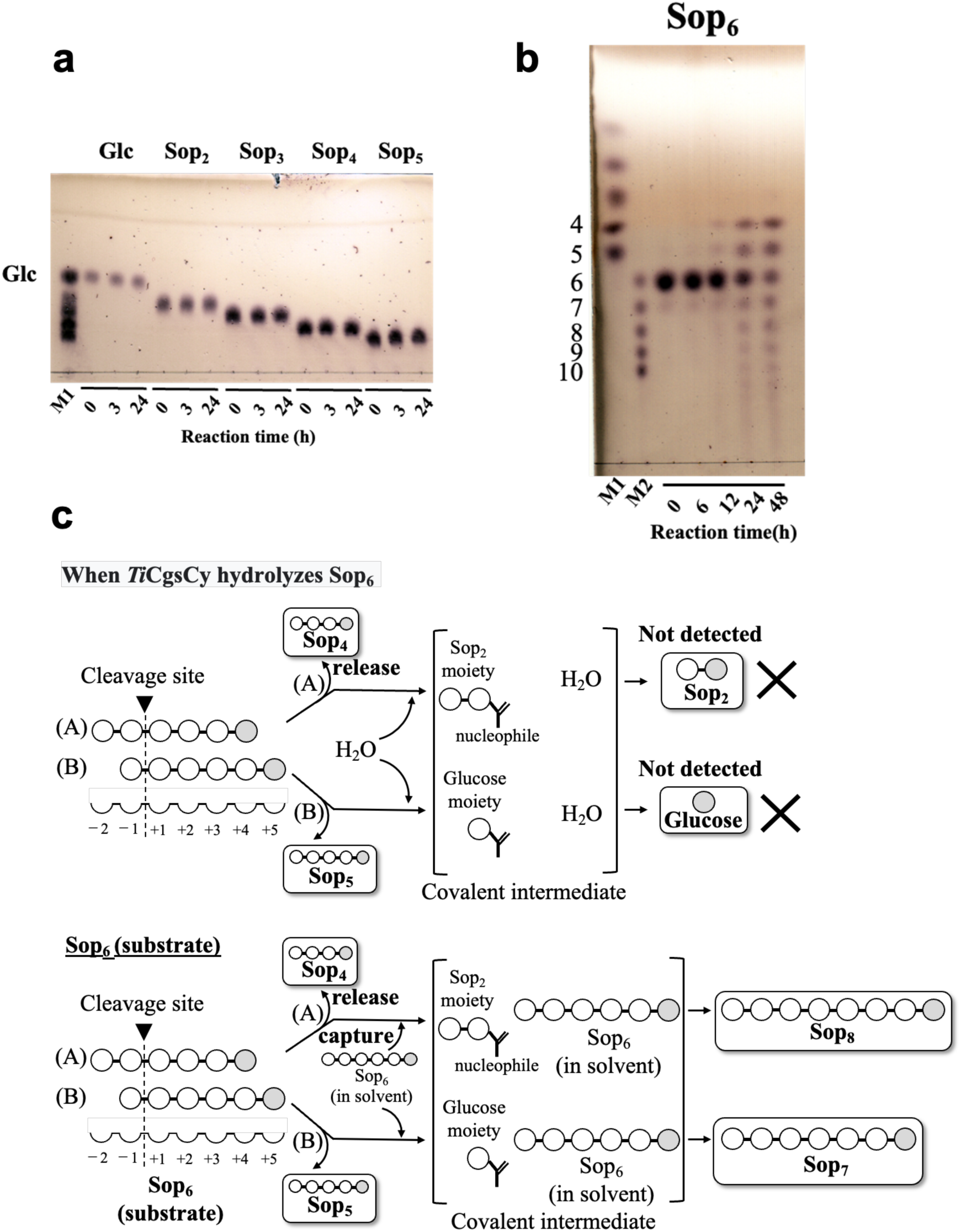
TLC analyses of activity toward Glc and Sop_2–6_. **a, b**, Lane M1, 5 mM glucose and Sop_2–5_. Lane M2, 5 mM Sop_6–10_. Each sample (0.2–1 μl) was spotted on the plate. **c**, Possible reaction steps of TiCGS_Cy_ with a substrate Sop_6_. Subsite binding is based on TLC results. The upper two reactions were not observed because H_2_O molecules does not participate in attacking covalent intermediates.

Generally, in the case of enzymes that produce cyclic glycan polymers, a nucleophile initially binds covalently to a substrate to form a glycosyl-enzyme intermediate. This intermediate is then subjected to nucleophilic attack either by a water molecule, an intermolecular hydroxy group, or an intramolecular hydroxy group to cause hydrolysis, disproportionation or cyclization reaction, respectively. In the case of TiCGS_Cy_ with Sop_6_, Sop_4_ and Sop_5_ were detected as a result of reaction, which suggested that Sop_6_ binds to TiCGS_Cy_ at subsite –2 to +4 or –1 to +5 (Figs 2b and 2c). However, glucose and Sop_2_ (counterparts of Sop_5_ and Sop_4_, respectively, when Sop_6_ is hydrolyzed) were not detected. This result indicates that TiCGS_Cy_ catalyzes only transglycosylation without hydrolysis (Fig. 2c). Likewise, with Sop_7–10_ as substrates, glucose and Sop_2–3_ were not detected (Fig. 3). Therefore, TiCGS_Cy_ requires at least four subsites (from subsite +1 to subsite +4) occupied by glucose moieties for the reaction to proceed.

**Figure 3.**
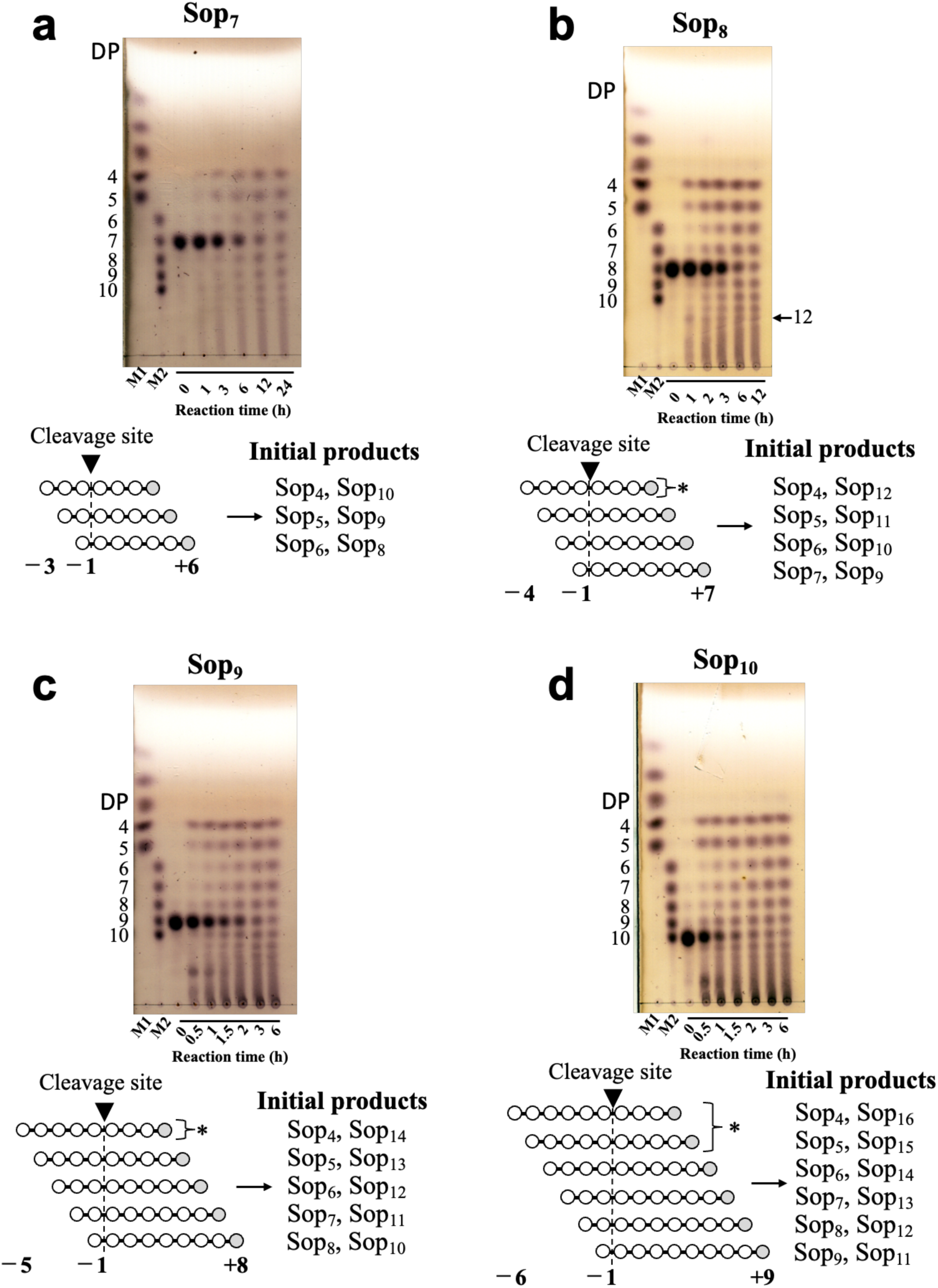
TLC analysis of activity toward Sop_7–10_. Lane M2, 5 mM glucose and Sop_2–5_. Lane M2, 5 mM Sop_6–10_. Asterisks represent preferential reaction patterns at the initial stage of the reactions.

In terms of the reaction velocities of substrates examined, the larger the DP of the substrates are, the faster the amount of the products reached to the similar level (Figs. 2 and 3). This result suggests that TiCGS_Cy_ prefers longer substrates, which is consistent with the biochemical property of CGS known to produce CβGs with DPs around 20. The amount of Sop_4–6_ produced from Sop_7_, Sop_8_ and Sop_9_ were Sop_4_ > Sop_5_ > Sop_6_, while in the case of Sop_10_ as a substrate, it was Sop_4–5_ > Sop_6_ (Fig. 3). These results suggest that subsites up to −5 are involved in substrate recognition.

### Overall structure of TiCGS_Cy_

A ligand-free structure of TiCGS_Cy_ was determined with 3.9 Å resolution (Table S1). An asymmetric unit contains almost identical four molecules (RMSD, 0.3 Å). The enzyme consists of a single (α/α)_6_-barrel domain with several inserted α-helices (Fig. 4ac). According to DALI server (Holm 2020), CpSGL (RMSD, 2.4 Å; sequence identity, 17%; PDB ID, 5GZH), GH144 enzyme from *Parabacteroides distasonis* (RMSD, 2.4 Å; sequence identity, 18%; PDB ID, 5Z06) and TfSGL (RMSD, 2.7 Å; sequence identity, 12%; PDB ID, 6IMU) came up as top 3 structurally similar proteins in the case TiCGS_Cy_ is set as a query structure. TiCGS_Cy_ is structurally similar to these three enzymes although amino acid sequence identities are very low. Structure-based multiple amino acid alignment suggests that the additional α-helices in the middle region is unique to CGSs, and they were not found in SGLs (Fig. 4c). A large pocket observed in (α/α)_6_-barrel domain is expected to be a substrate binding site of TiCGS_Cy_ (Fig. 4b).

**Figure 4.**
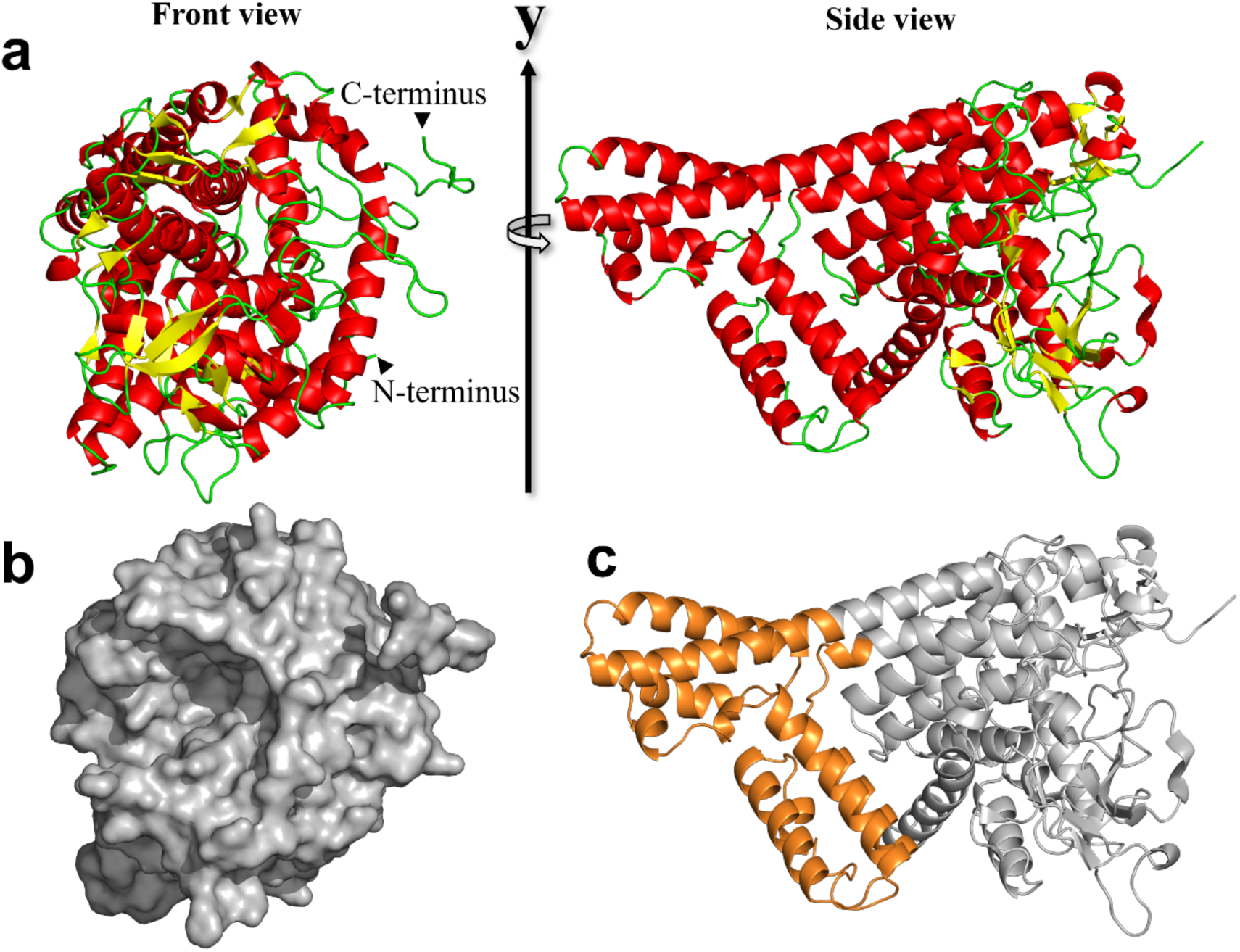
Overall structure of TiCGS_Cy_. Cartoon (**a, c**) and surface (**b**) representations of TiCGS_Cy_. α-Helices and β-strands are shown in red and yellow, respectively. The surface is shown in gray. The additional α-helices domain observed in TiCGS_Cy_ is shown in orange.

### Comparison of substrate-binding site of TiCGS_Cy_ with CpSGL and TfSGL

To analyze a substrate binding mode of TiCGS_Cy_, TiCGS_Cy_ was superimposed with two enzymes: TfSGL complexed with a substrate (Sop_7_) and CpSGL complexed with a glucose and a Sop_3_. Consequently, the three overall structures are superimposed well (Fig. S5). A shape of the substrate pocket of TiCGS_Cy_ is similar to those of TfSGL and CpSGL in that the substrates observed in TfSGL and CpSGL complex structures can be potentially accommodated in the pocket, although the superimposed glucose moiety at subsite −3 is a little too close to TiCGS_Cy_ (Fig. 5a). There is a sufficient space beyond 2-OH group of the potential subsite −4 (Fig. 5b), which is consistent with the fact that TiCGS_Cy_ prefers Sop_n_s with larger DPs (Fig. 4). Contrarily, W1394 is likely to block binding of glucose moieties beyond subsite +3 (Fig. 5c). Taking into account the result of action pattern analysis that subsite +4 is necessary (Figs. 3 and 4), side chain of W1394 may flip to make room for substrate binding. Overall, it is suggested that the pocket in the (α/α)_6_-barrel domain is the substrate binding site.

**Figure 5.**
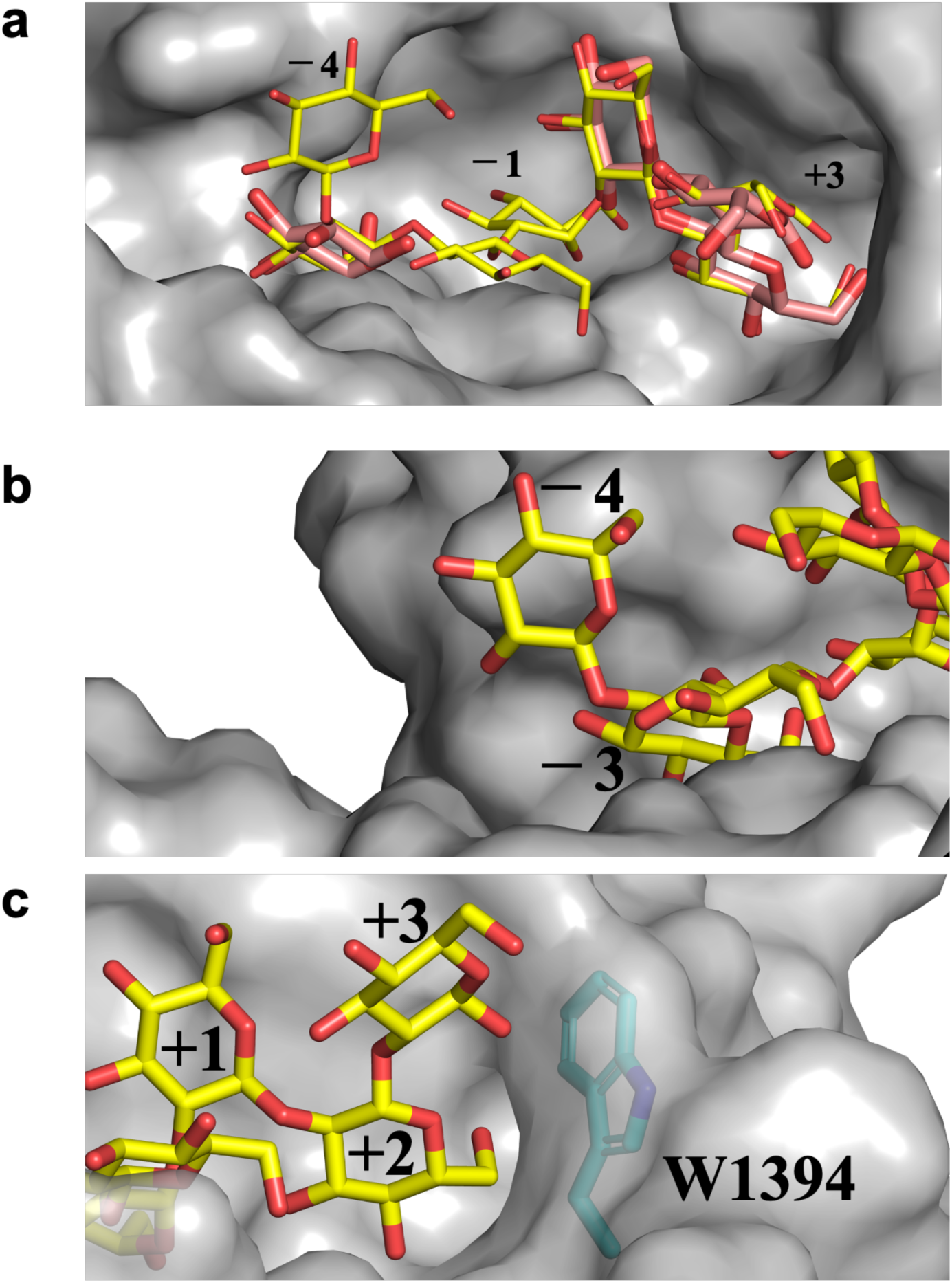
Substrate pocket of TiCGS_Cy_. The substrate pocket of TiCGS_Cy_ is shown semi-transparently in gray. Sop_7_ molecules shown as yellow sticks are placed by superimposition of TfSGL-Sop_7_ complex structure. Glucose and Sop_3_ molecules shown as light red sticks are placed by superimposition of CpSGL-glucose, Sop_3_ complex structure. Number labels represent subsite positions. **b**, **c** Close-up views around subsites −4 (**b**) and +3 (**c**).

### Catalytic residues of TiCGS_Cy_

Canonical enzymes that synthesize cyclic carbohydrates take advantage of anomer-retaining mechanism to achieve transglycosylation reaction (See https://www.cazypedia.org/index.php/Transglycosylases for details) (Sinnott 1990). First, a nucleophilic acidic residue attacks an anomeric carbon atom at subsite −1 to form a glycosyl-enzyme covalently bonded intermediate, and an acid/base catalyst provides a proton to a scissile bond oxygen atom in a substrate to release a moiety at the reducing end from the scissile bond. This step is called glycosylation step. In the next step called deglycosylation, the intermediate is attacked by an intramolecular hydroxy group to complete a cyclization reaction mediated by an acid/base catalyst.

In the case of TiCGS_Cy_, E1442 was found with a clear electron density at the position corresponding to that of the nucleophilic water in TfSGL, which attacks the anomeric carbon of the glucose moiety at subsite −1 (Fig. 6ab). Meanwhile, no candidate acidic residue directly interacting with a scissile bond oxygen atom is found. However, E1356 of TiCGS_Cy_ is well-superimposed with E262 of TfSGL, a clearly evidenced catalytic residue acting on a scissile bond of a substrate through 3-OH of a glucose moiety at subsite +2 (Fig. 6a). Electron density of E1356 was also observed clearly (Fig. 6b). Both E1442Q and E1442A mutants showed no cyclization activity toward LβGs, and E1356A mutant showed drastically reduced cyclization activity toward LβG (Fig. S6). These results strongly suggest that E1442 is a catalytic residue that acts as a nucleophile, and E1356 is an acid/base catalyst.

**Figure 6.**
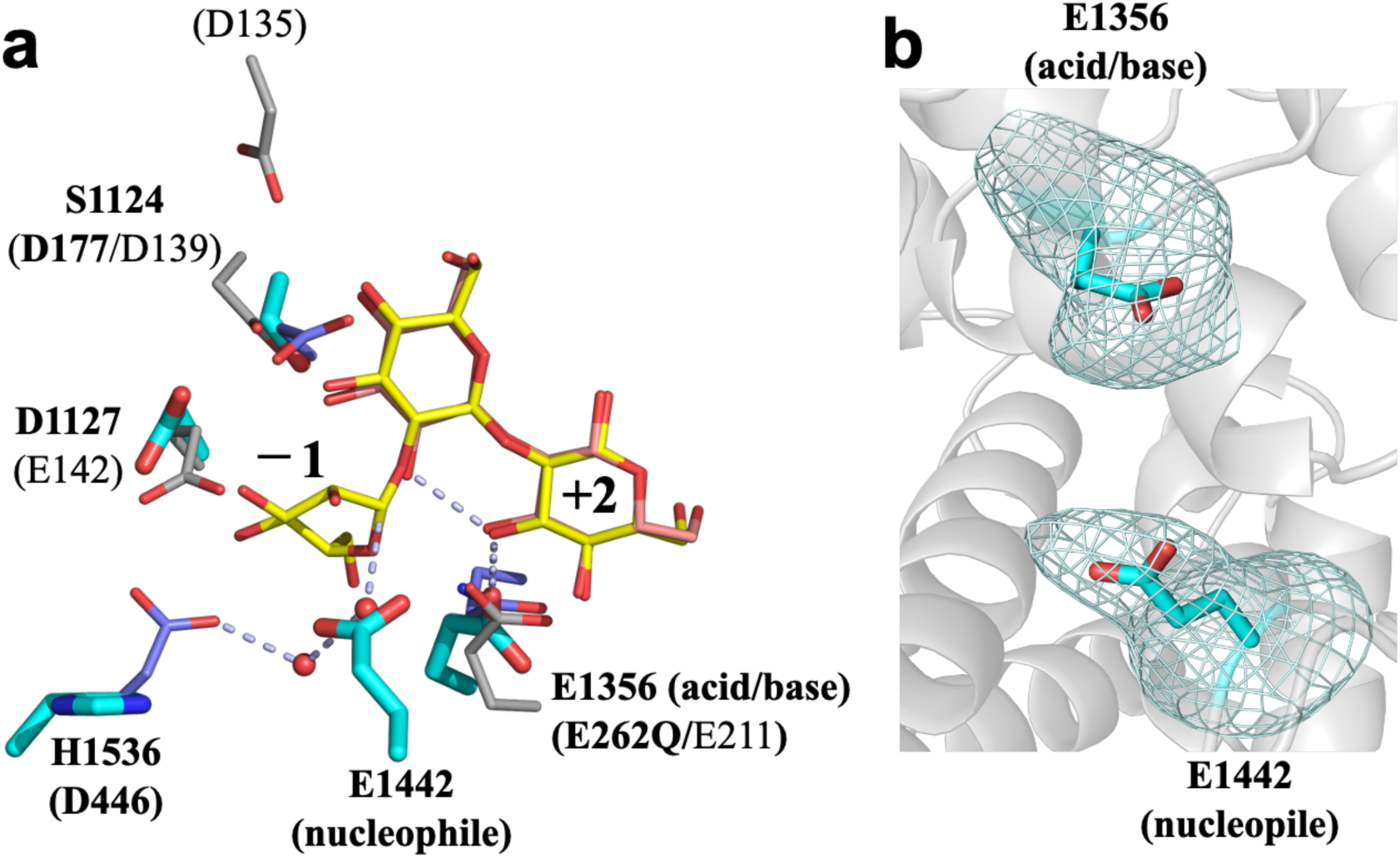
Superimposition of catalytic residues and related residues in TiCGS_Cy_, TfSGL and CpSGL. **a**, Sop_7_ in TfSGL-Sop_7_ complex and Sop_3_ in CpSGL in CpSGL-Glc, and Sop_3_ complex are partially visualized as yellow and light red sticks, respectively. Residues in TiCGS_Cy_, TfSGL and CpSGL are shown as thick cyan, purple and gray sticks, respectively. Residues in TiCGS_Cy_, TfSGL and CpSGL are labelled with bold letters, bold letters in parentheses and plane letters in parentheses, respectively. Water molecules observed in TfSGL-Sop_7_ complex are shown as red spheres. Gray dashed lines represent a route of the reaction pathway in TfSGL. Subsite positions are labelled −1 and +2. **b**, A *F*_o_-*F*_c_ omit map for E1356 and E1442 in TiCGS_Cy_. The map is shown at the 4.0σ contour level and represented as cyan meshes.

According to prediction of p*K*_a_ by PROPKA3.5.0 (Olsson et al. 2011), p*K*_a_ values of E1442 and E1356 in chain A were 6.72 and 8.98, respectively. E1400 highly conserved among CGSs is found in the vicinity of E1442 although E1400 is not conserved in TfSGL or CpSGL (Fig. 7). A negative charge of E1400 is considered to raise the p*K*_a_ value of E1442. Contrarily, the basic residues H1536 and H1537 are found in close proximity to E1356, which probably contribute to the decrease in p*K*_a_ value of E1356. In addition, these two histidine residues are also highly conserved among CGSs (Fig. S1). The difference of the p*K*_a_ values between E1442 and E1356 suggests that these two residues are catalysts.

**Figure 7.**
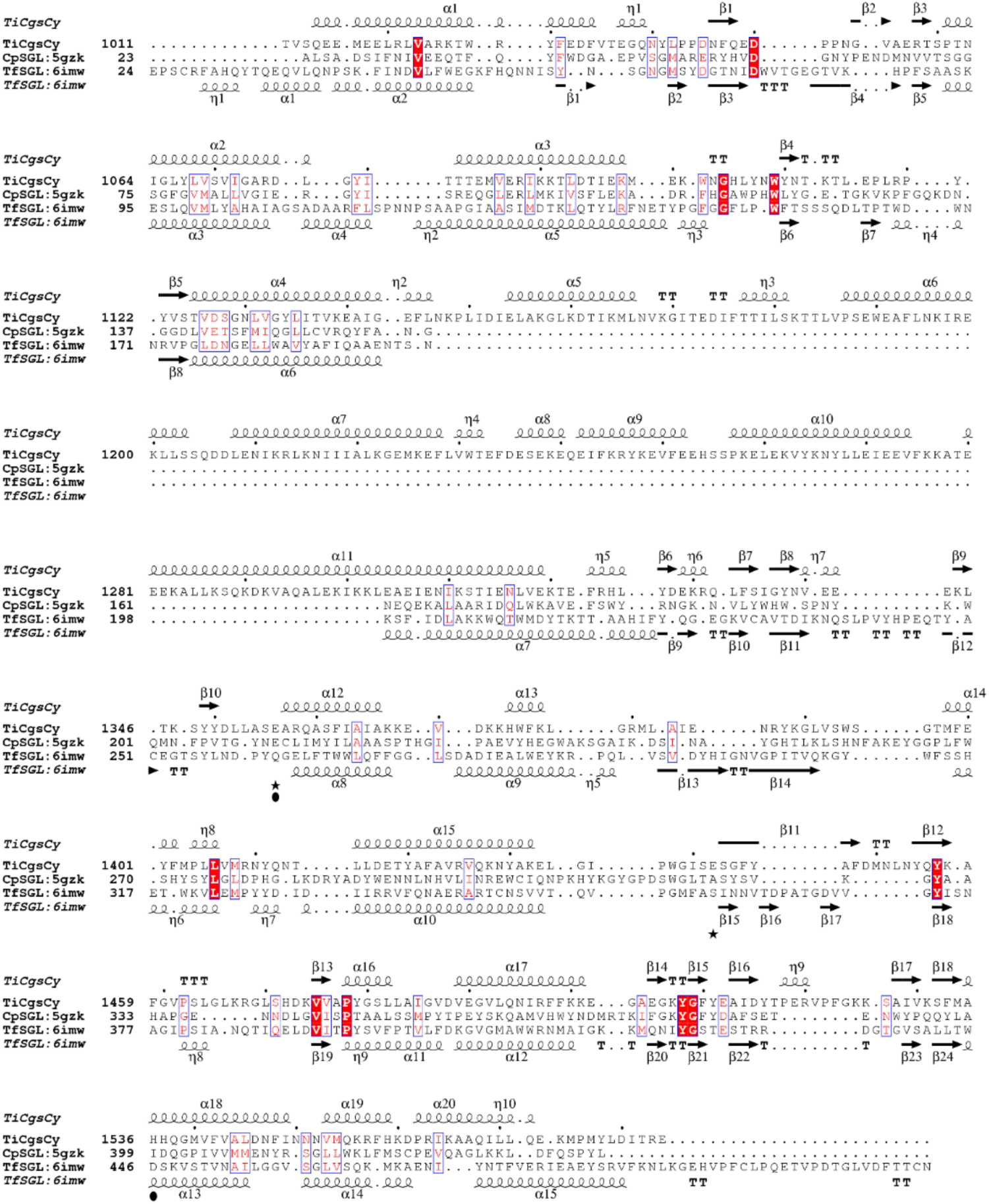
Multiple amino acid alignment of TiCGS_Cy_, TfSGL and CpSGL. Closed stars and circles represent catalytic residues of TiCGS_Cy_ and TfSGL, respectively.

## Discussion

In the present study, we explicitly showed that TiCGS_Cy_ domain alone produced CβGs by transglycosylation reaction without hydrolysis. Preference of this domain for Sop_n_s with higher DPs is consistent with the chain lengths of reaction products by the intact CGSs (Hisamatsu 1992; Ciocchini et al. 2007; Guidolin et al. 2009). Recently, a cryo-EM structure of intact CGS from *A. tumefaciens* has been reported (Sedzicki *et al*. 2023). Nevertheless, a plausible reaction pathway to account for transglycosylation could not be drawn from the structure because the orientation of a substrate chain shown in the manuscript appears to be reversed in comparison with that determined in TfSGL. Reaction products from LβGs using the intact CGS have not been identified as CβGs, and the possibility of transglycosylation led by reversible phosphorolysis with a GH94 glycoside phosphorylase domain cannot be excluded. Thus, the present study is indeed the first demonstration of detailed reaction mechanism of CGS_Cy_ domain.

Based on the overall results of structural and functional analysis of TiCGS_Cy_, the reaction pathway of the enzyme can be explained as follows (Figs. 6 and 8). First, in the glycosylation step, E1356 acts as a general acid to provide a proton to a scissile bond oxygen atom through 3-OH group of a glucose moiety at subsite +2. Simultaneously, E1442 attacks an anomeric center at subsite −1 as a nucleophile to form a glycosyl-enzyme intermediate. Next, E1356 acts as a general base to draw a proton of inter- or intra-molecular 2-OH group of a glucose moiety at subsite +2 in a reversed manner. The activated (deprotonated) 2-OH group attacks the anomeric carbon of covalently bonded glucose moiety to release transglycosylation products. If an intermolecular hydroxy group attacks an anomeric carbon atom, a product in a linear form is released. In the case of an intramolecular hydroxy group, a product in a cyclic form is released (Fig. 8). Although such proton transfer called Grotthuss mechanism is non-canonical among GH families (de Grotthuss 1806; Cukierman 2006), GH162 and GH186 SGLs (TfSGL and OpgD from *E. coli*, respectively) (Tanaka *et al*. 2019; Motouchi *et al*. 2023), GH130 4-*O*-β-D-mannosyl-D-glucose phosphorylase (Nakae et al. 2013) and GH136 lacto-*N*-biosidase (Yamada et al. 2017) are isolated examples of Grotthuss mechanism, supporting the proposed reaction mechanism of TiCGS_Cy_.

**Figure 8.**
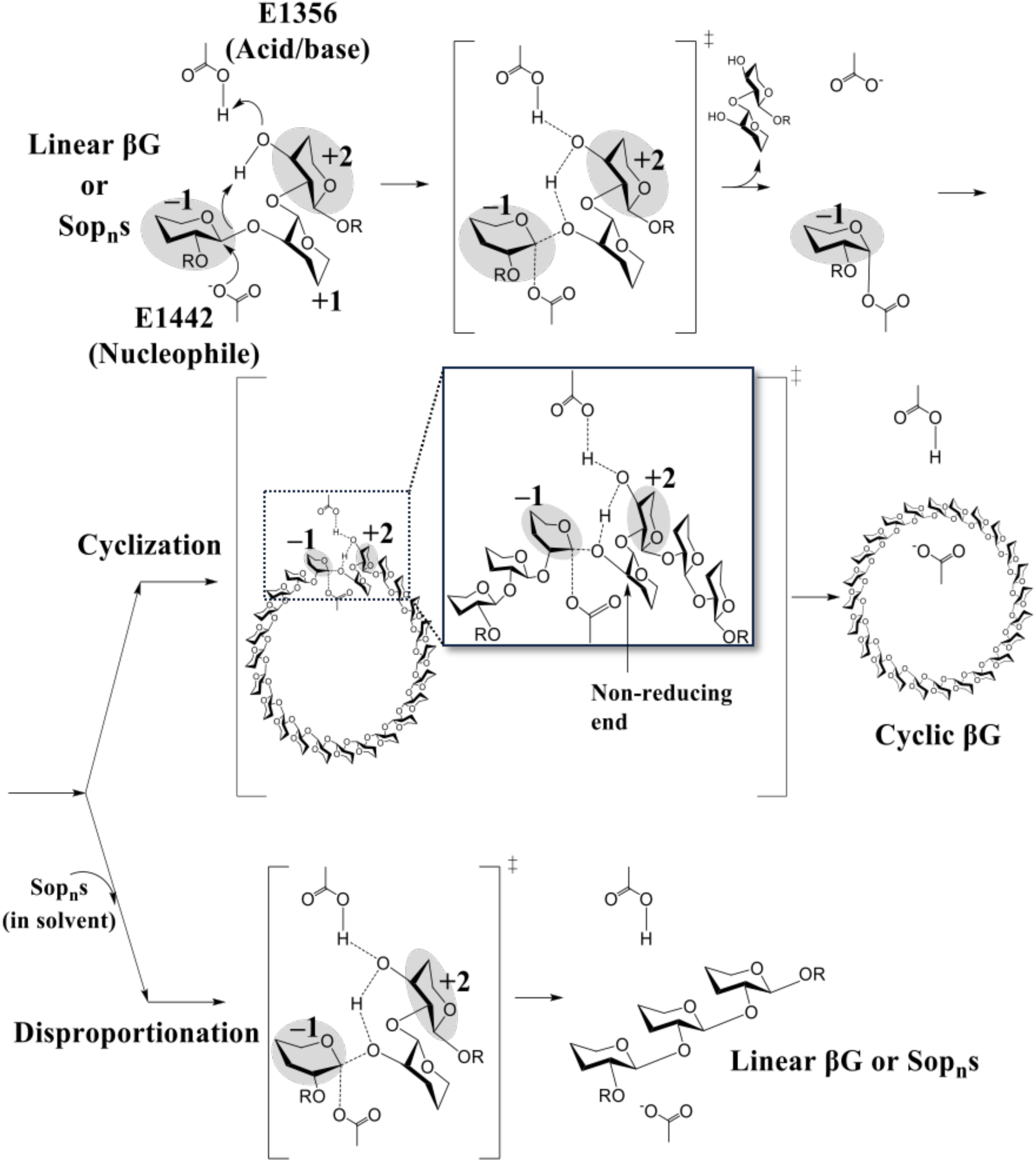
Schematic representation of the proposed reaction mechanism of TiCGS_Cy_.

Comparison of the reaction mechanisms between GH144, GH162 and CGS revealed that the general acid (E262 in GH162 TfSGL) is shared with GH144 CpSGL (E211) and TiCGS_Cy_ (E1356, in anomer-retaining mechanism, an acid/base) (Fig. 6). These residues are also conserved in multiple amino acid alignment (Fig. 7). Contrarily, D446 in TfSGL (a general base) is substituted in CpSGL with a hydrophobic residue that cannot act as a catalyst (Fig. 6), indicating that the reaction pathways of GH162 and GH144 are different although the reaction pathway of GH144 has not been fully determined (Tanaka et al. 2019). In TiCGS_Cy_, D446 of TfSGL is substituted with H1536 (Fig. 6). This histidine is a proton dissociative and highly conserved residue among CGSs (Fig. S1). Nevertheless, E1442 is a nucleophile and H1536 is considered to play an important role in supporting deprotonation of E1442. This observation clearly indicates the difference in the reaction pathways between GH162 and CGSs. Although GH162 TfSGL, GH144 CpSGL and TiCGS_Cy_ share a similar overall fold, they belong to phylogenetically far different groups. Considering the clear difference in the reaction mechanism between the three groups, we propose that the group of CGSs including TiCGS be given a new GH family, GHxxx. This study provides significant insights into biosynthesis of the physiologically important CβGs by further understanding of structures and functions of CGSs. Moreover, this finding is a large achievement to expand the field of carbohydrate-active enzymes by adding a new group of enzymes.

## Acknowledgments

This work was supported by Photon Factory for X-ray data collection (Proposal No. 2020G527 and No. 2021G685). The CβG with DP of 17–24 was kindly provided by Dr. Hisamatsu (Mie University).

## Author Contribution Statement

MN, HN, HT and TM conceived and designed research. NT, RS, KKo, HN, SK, KKu and MN conducted experiments. NT, HN and MN contributed to preparation of compounds for analysis. NT, RS, MN and TM analyzed the data. NT, MN and TM wrote the draft manuscript. All the authors read and approved the content of the manuscript.

