## Supplemental_Fig_Table for "Functional and structural analysis of a cyclization domain in a cyclic β-1,2-glucan synthase"

**Table S1 Data collection and refinement statistics**

| Data set | $TiCGS_{Cy}$ |
|---|---|
| Beamline | BL-5A |
| Space group | $P4_12_12$ |
| **Data collection** | |
| Unit cell parameters (Å) | $a$ = 172.72, $b$ = 172.72, $c$ = 395.60 |
| Resolution (Å)[a] | 89.60–3.90 (4.01–3.90) |
| Total reflections[a] | 1,385,923 (116,994) |
| Unique reflections[a] | 55,512 (4,497) |
| Completeness (%)[a] | 100.0 (99.9) |
| Redundancy[a] | 25.0 (26.0) |
| Mean $I/\sigma$ ($I$)[a] | 9.8 (2.9) |
| $R_{merge}$ (%)[a] | 0.47 (1.79) |
| $R_{pim}$ (%)[a] | 0.49 (1.85) |
| $CC_{1/2}$[a] | 0.99 (0.85) |
| **Refinement** | |
| Resolution (Å) | 89.60–3.90 |
| No. of unique reflections | 52,696 |
| Completeness (%)[a] | 99.96 |
| $R_{work}/R_{free}$ (%) | 19.8/23.2 |
| **PDB entry** | xxxx |

[a] Values in parentheses represent the highest resolution shell.

**Table S2 Primers used in this study**

| Preparation of $TiCGS_{Cy}$ mutants (5'-oligonucleotide-3') | | |
|---|---|---|
| **Mutants** | **Forward** | **Reverse** |
| E1442A | ATATCAGCGTCGGGATTTTACGCTTTT | TCCCGACGCTGATATGCCCCACGGAAT |
| E1442Q | ATATCACAGTCGGGATTTTACGCTTTT | TCCCGACTGTGATATGCCCCACGGAAT |
| E1356A | GCTTCAGCAGCAAGGCAAGCGAGTTTT | CCTTGCTGCTGAAGCCAACAAGTCATA |

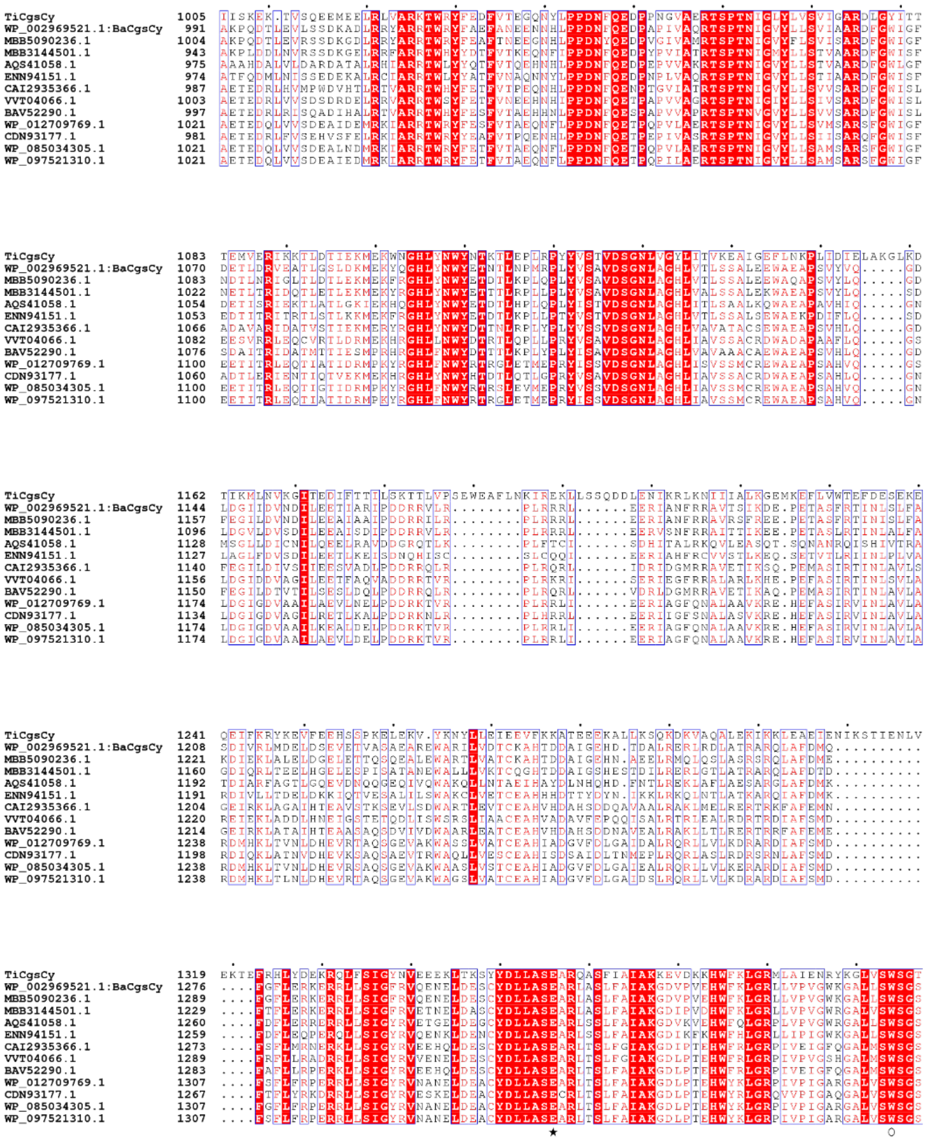
TiCgsCy
WP_002969521.1:BaCgsCy
MBB5090236.1
MBB3144501.1
AQS41058.1
ENN94151.1
CAI2935366.1
VVT04066.1
BAV52290.1
WP_012709769.1
CDN93177.1
WP_085034305.1
WP_097521310.1

continued

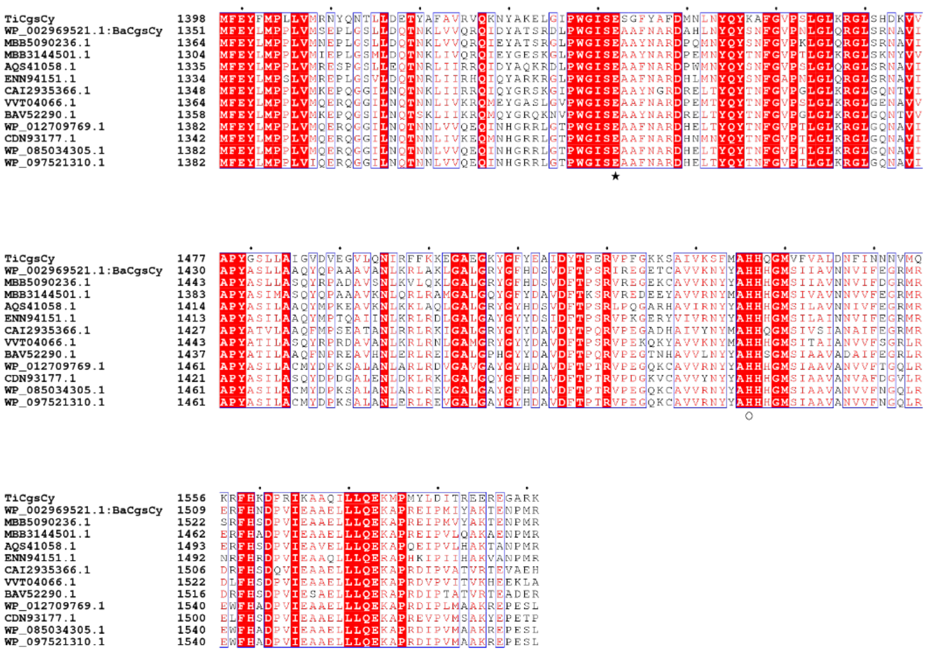

**Figure S1 Multiple amino acid alignment of CGSs**

Only cyclization domain region is shown. Closed stars indicate catalytic residues of $TiCGS_{Cy}$. W1394 and H1536 are labelled with open circles.

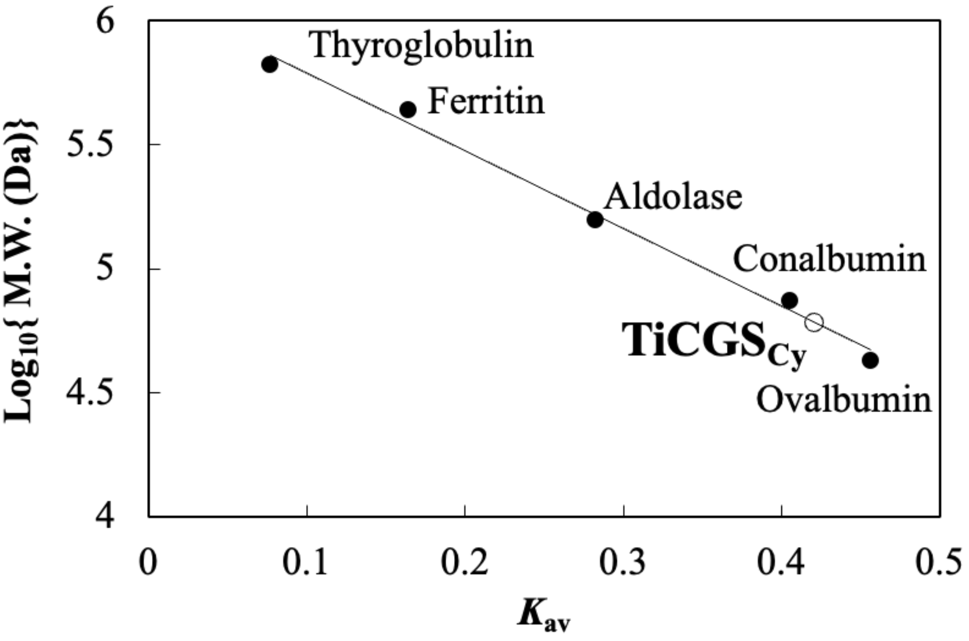


**Figure S2 Size-exclusion chromatography of TiCGS$_{Cy}$**

Molecular weight protein markers and TiCGS$_{Cy}$ are shown in closed and open circles, respectively.

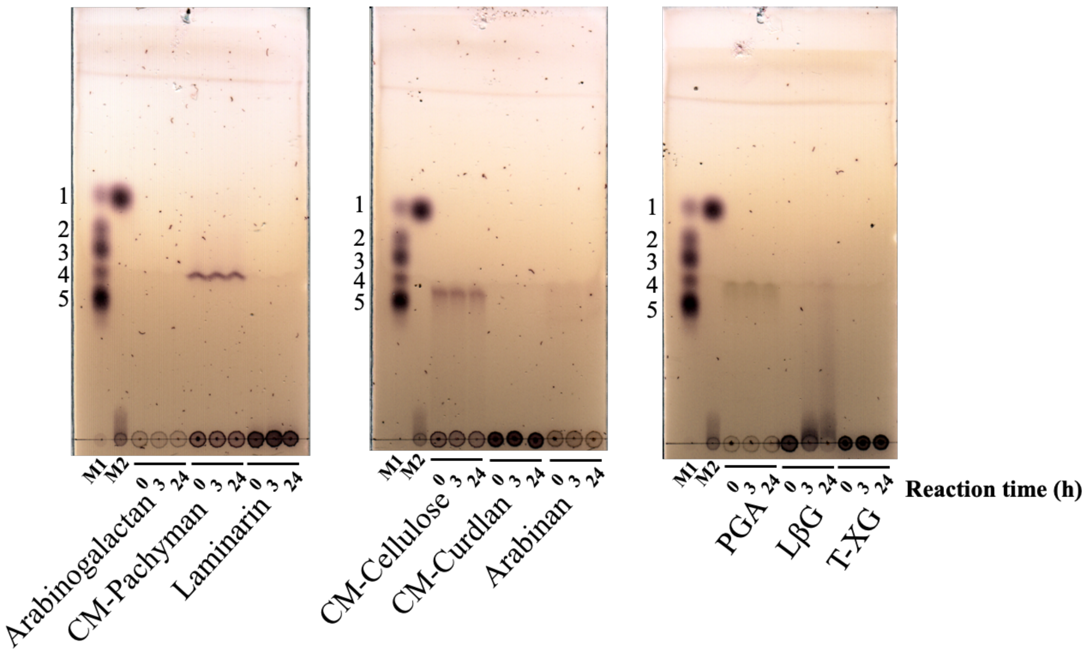


**Figure S3 Substrate specificity of TiCGS$_{Cy}$**

Lane M1, 0.5 µl of mixture of glucose (5 mM) and Sop$_{2–5}$ (each 5 mM) was spotted. Lane M2, a sample for NMR after sufficient BGL-treatment. Numbers beside the TLC plates represent DPs of Glc and Sop$_{2–5}$.

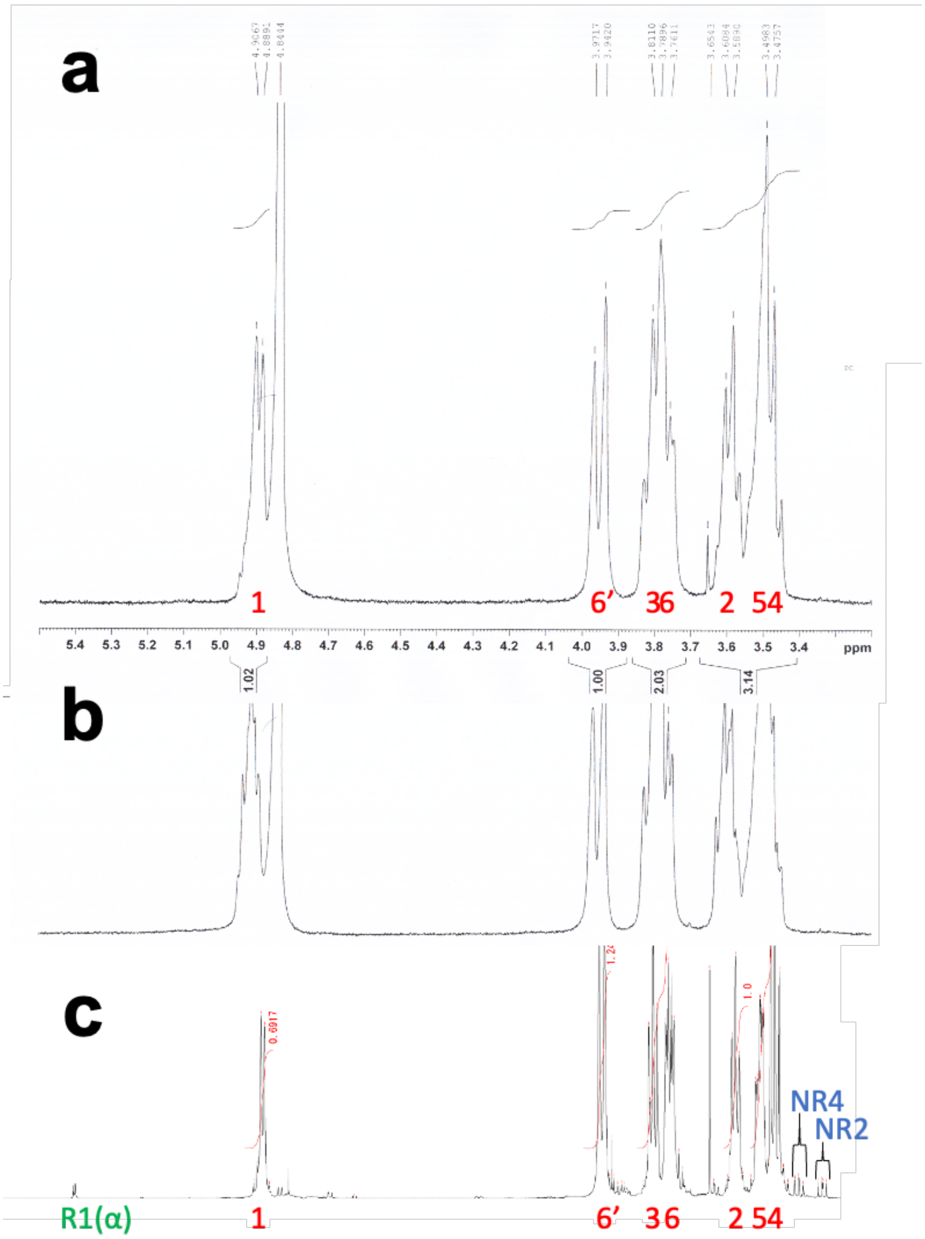


**Figure S4 [1]H-NMR of the BGL-resistant reaction products**

**a**, The BGL-resistant reaction products from LβG catalyzed by $TiCGS_{Cy}$. **b**, CβG with DP17–24 provided by Dr. Hisamatsu (Hisamatus et al., 1984). **c**, LβG with average DP of 39 produced by SOGP from *L. innocua* (Nakajima et al., 2014). Colored numbers represent positions of protons covalently bonded to carbon atoms. Red, blue and green letters represent inner, non-reducing end and reducing end glucose moieties, respectively. (α) represents α-anomer glucans.

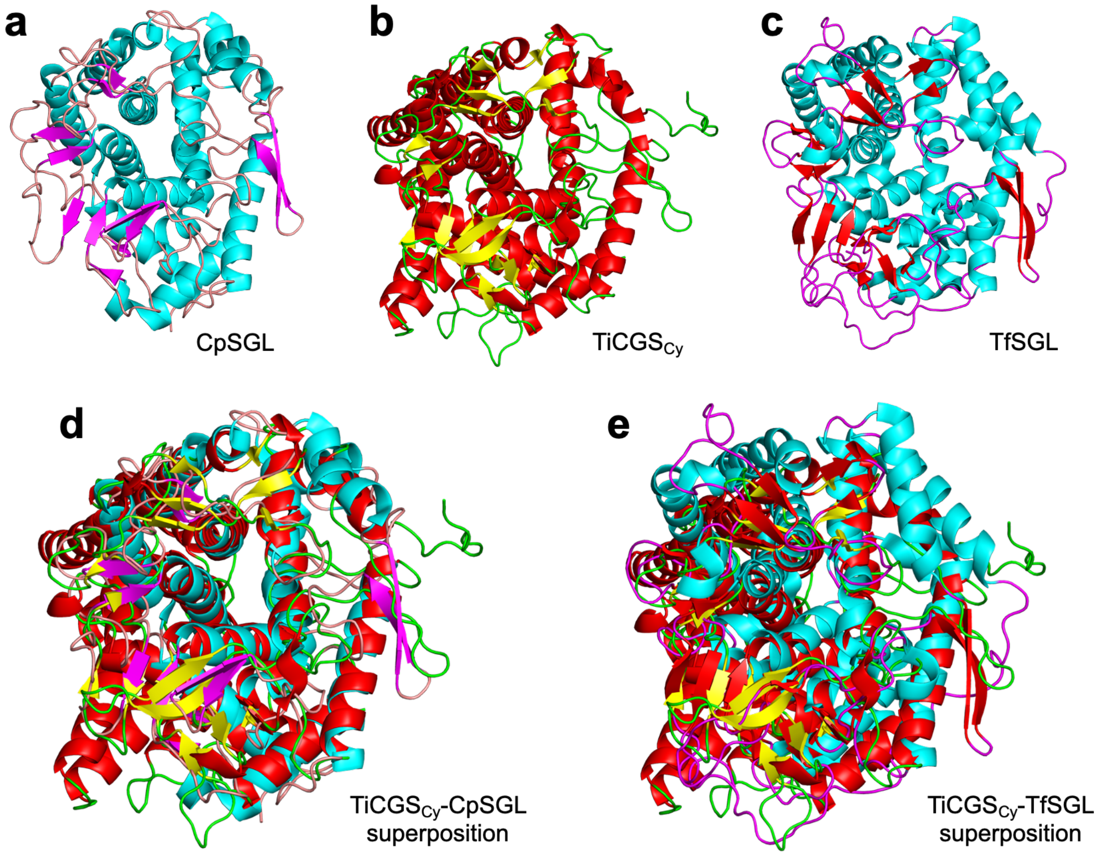


**Figure S5 Superimposition of TiCGS$_{Cy}$ with CpSGL, and TfSGL.**

The three enzymes are shown in cartoon and are colored by secondary structures.

**a**, **b**, **c**, Sole structures of CpSGL, TiCGScy, and TfSGL respectively. **d**, Superimposition of TiCGScy with CpSGL. **e**, Superimposition of TiCGScy with TfSGL.

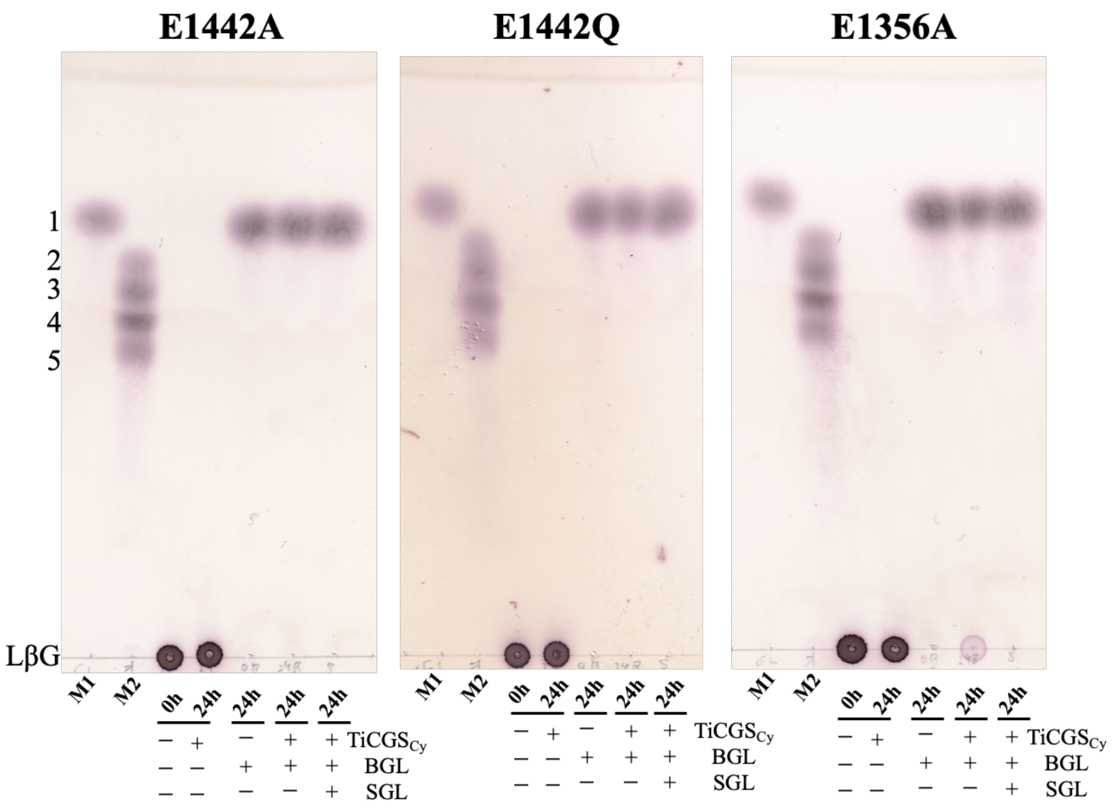


**Figure S6 TLC analyses of mutant TiCGS$_{Cy}$s**

Lane M1, glucose (5 mM, 0.5 μl) was spotted. Lane M2, a 0.5 μl of mixture of Sop$_{2–5}$ (each mM) was spotted. Horizontal lines on the TLC plates represent origins. Each sample (0.5–2 μl) was spotted on the origin.
